# Using high-resolution variant frequencies to empower clinical genome interpretation

**DOI:** 10.1101/073114

**Authors:** Nicola Whiffin, Eric Minikel, Roddy Walsh, Anne O’Donnell-Luria, Konrad Karczewski, Alexander Y Ing, Paul JR Barton, Birgit Funke, Stuart A Cook, Daniel MacArthur, James S Ware

## Abstract

Whole exome and genome sequencing have transformed the discovery of genetic variants that cause human Mendelian disease, but discriminating pathogenic from benign variants remains a daunting challenge. Rarity is recognised as a necessary, although not sufficient, criterion for pathogenicity, but frequency cutoffs used in Mendelian analysis are often arbitrary and overly lenient. Recent very large reference datasets, such as the Exome Aggregation Consortium (ExAC), provide an unprecedented opportunity to obtain robust frequency estimates even for very rare variants. Here we present a statistical framework for the frequency-based filtering of candidate disease-causing variants, accounting for disease prevalence, genetic and allelic heterogeneity, inheritance mode, penetrance, and sampling variance in reference datasets. Using the example of cardiomyopathy, we show that our approach reduces by two-thirds the number of candidate variants under consideration in the average exome, and identifies 43 variants previously reported as pathogenic that can now be reclassified. We present precomputed allele frequency cutoffs for all variants in the ExAC dataset.

## INTRODUCTION

Whole exome and whole genome sequencing have been instrumental in identifying causal variants in Mendelian disease patients^1^. As every individual harbors ~12,000-14,000 predicted protein-altering variants^2^, distinguishing disease-causing variants from benign bystanders is perhaps the principal challenge in contemporary clinical genetics. A variant’s low frequency in, or absence from, reference databases is now recognised as a necessary, but not sufficient, criterion for variant pathogenicity^3,4^. The recent availability of very large reference databases, such as the Exome Aggregation Consortium (ExAC)^2^ dataset, which has characterised the population allele frequencies of 10 million genomic variants through the analysis of exome sequencing data from over humans, provides an opportunity to obtain robust frequency estimates even for rare variants, improving the theoretical power for allele frequency filtering in Mendelian variant discovery efforts.

In practice, there exists considerable ambiguity around what allele frequency should be considered “too common”, with the lenient values of 1% and 0.1% often invoked as conservative frequency cutoffs for recessive and dominant diseases respectively^5^. Population genetics, however, dictates that severe disease-causing variants must be much rarer than these cutoffs, except in cases of bottlenecked populations, balancing selection, or other special circumstances^6,7^.

It is intuitive that when assessing a variant for a causative role in a dominant Mendelian disease, the frequency of a variant in a reference sample, not selected for the condition, should not exceed the prevalence of the condition^8,9^. This rule must, however, be refined to account for different inheritance modes, genetic and allelic heterogeneity, and reduced penetrance. In addition, for rare variants, estimation of true population allele frequency is clouded by considerable sampling variance, even in the largest samples currently available. These limitations have encouraged the adoption of very lenient approaches when filtering variants by allele frequency^10,11^, and recognition that more stringent approaches that account for disease-specific genetic architecture are urgently needed^8^.

Here we present a statistical framework for assessing whether rare variants are sufficiently rare to cause penetrant Mendelian disease, while accounting for both architecture and sampling variance in observed allele counts. We demonstrate that allele frequency cutoffs well below 0.1% are justified for a variety of human disease phenotypes and that such filters can remove an additional two-thirds of variants from consideration when compared to traditionally lenient frequency cutoffs. We present precomputed allele frequency filtering values for all variants in the Exome Aggregation Consortium database, which are now available through the ExAC data browser and for download, to assist others in applying our framework.

## RESULT

### Defining the statistical framework

For a penetrant dominant Mendelian allele to be disease causing, it cannot be present in the general population more frequently that the disease it causes. Furthermore, if the disease is genetically heterogeneous, it must not be more frequent than the proportion of cases attributable to that gene, or indeed to any single variant. We can therefore define the maximum credible population allele frequency (for a pathogenic allele) as:

*maximum credible population AF = prevalence × maximum allelic contribution × 1/penetrance*

where *maximum allelic contribution* is the maximum proportion of cases potentially attributable to a single allele, a measure of heterogeneity.

We do not know the true population allele frequency of any variant, having only an observed allele frequency in a finite population sample. Moreover, confidence intervals around this observed frequency are problematic to estimate given our incomplete knowledge of the frequency spectrum of rare variants, which appears to be skewed towards very rare variants. For instance, a variant observed only once in a sample of 10,000 chromosomes is much more likely to have a frequency < 1:10,000 than a frequency >1:10,000.^2^

If we turn the problem around, and begin instead from allele frequency, specifying a maximum *true* allele frequency value we are willing to consider in the population (using the equation above), then we can estimate the probability distribution for allele counts in a given sample size. This follows a binomial distribution, and can be satisfactorily approximated with a Poisson distribution (see **Online Methods**). This allows us to set an upper limit on the number of alleles in a sample that is consistent with a given population frequency.

Taking a range of cardiac disorders as exemplars, we use this framework to define the maximum credible allele frequency for disease-causing variants in each condition, and define and validate a set of maximum tolerated allele counts in the ExAC reference population sample. Figure 1 shows the general outline of our approach.

**Figure 1.**
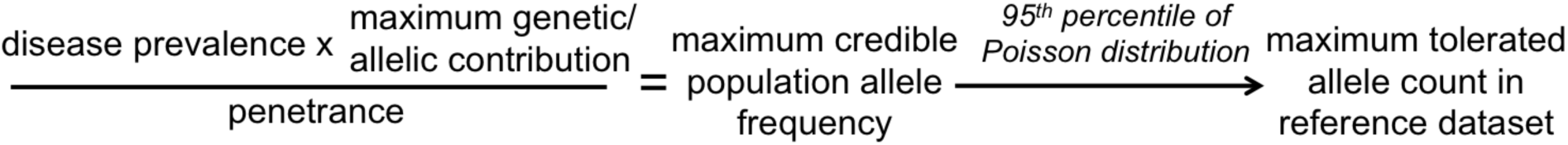
A flow diagram of our approach, applied to a dominant condition. First, a disease-level maximum credible population allele frequency is calculated, based on disease prevalence, heterogeneity and penetrance. This is then used to calculate the maximum tolerated allele count in a reference dataset, taking into acount the size of this dataset.

### Application and validation in hypertrophic cardiomyopathy

We illustrate our general approach using the dominant cardiac disorder hypertrophic cardiomyopathy (HCM), which has an estimated prevalence of 1 in 500 in the general population^12^. As there have been previous large-scale genetic studies of HCM, with series of up to 6,179 individuals^12,13^, we can make the assumption that no newly identified variant will be more frequent in cases that those identified to date (at least for well-studied ancestries). This allows us to define the maximum allelic contribution of any single variant to the disorder. In these large case series, the largest proportion of cases is attributable to the missense variant *MYBPC3* c.1504C>T (p.Arg502Trp), found in 104/6179 HCM cases (1.7%; 95%CI 1.4-2.0%)^12,13^. We therefore take the upper bound of this proportion (0.02) as an estimate of the maximum allelic contribution in HCM (Table 1). Our maximum expected population allele frequency for this allele, assuming 50% penetrance as previously reported^14^, is 1/500 × 1/2 (dividing prevalence per individual by the number of chromosomes per individual) x 0.02 x 1/0.5 = 4.0x10^-5^, which we take as the maximum credible population AF for any causative variant for HCM (Table 1).

**Table 1.** Details of the most prevalent pathogenic variants in case cohorts for five cardiac conditions. Shown along with the frequency in cases is the estimated population allele frequency (calculated as: case frequency x disease prevalence x 1/2 x 1/variant penetrance) and the observed frequency in the ExAC dataset. *As penetrance estimates for individual variants are not widely available, we have applied an estimate of 0.5 across all disorders ( see **Supplementary information**). HCM k hypertrophic cardiomyopathy; DCM k dilated cardiomyopathy; ARVC k arrhythmogenic right ventricular cardiomyopathy; LQTS k long QT syndrome. Case cohorts and prevalence estimates were obtained from: HCM^12,13^, DCM^12,13,15^, ARVC^13,16^, LQTS^17,18^ and Brugada^19^,^20^.

| Disease | Prevalence | Commonest causative variant | Case count | Case frequency (95% CI) | Penetrance* | Expected population frequency (95%CI) | Model predicted maximum ExAC AC | Observed ExAC AC |
| --- | --- | --- | --- | --- | --- | --- | --- | --- |
| HCM | 1/500 | MYBPC3 | 104/6179 | 1.7% | 0.5 | 3.4x10<sup>-5</sup> | 9 | 3 |
|  |  | c.1504C>T |  | (1.4-2.0%) |  | (2.7-4.0x10<sup>-5</sup>) |  |  |
| DCM | 1/250 | TNNT2 | 18/1254 | 1.4% | 0.5 | 5.6x10<sup>-5</sup> | 16 | 0 |
|  |  | c.629_631delAGA |  | (0.78-2.1%) |  | (3.1-8.4x10<sup>-5</sup>) |  |  |
| ARVC | 1/1000 | PKP2 | 24/361 | 6.7% | 0.5 | 6.7x10<sup>-5</sup> | 17 | 6 |
|  |  | c.2146-1G>C |  | (4.1-9.2%) |  | (4.1-9.2x10<sup>-5</sup>) |  |  |
| LQTS | 1/2000 | KCNQ1 | 30/2500 | 1.2% | 0.5 | 6.0x10<sup>-6</sup> | 3 | 0 |
|  |  | c.797T>C |  | (0.77-1.6%) |  | (3.9-8.2x10<sup>-6</sup>) |  |  |
| Brugada | 1/1000 | SCN5A | 14/2111 | 0.66% | 0.5 | 6.6x10<sup>-6</sup> | 3 | 0 |
|  |  | c.5350G>A |  | (0.32-1.0%) |  | (0.32-1.0x10<sup>-5</sup>) |  |  |

To apply this threshold while remaining robust to chance variation in observed allele counts, we ask how many times a variant with population allele frequency of 4.0x10^-5^ can be observed in a random population sample of a given size. For a 5% error rate we take the 95th percentile of a poisson distribution with λ = expected allele count, which is given by: sample size (chromosomes) x expected population allele frequency (**Online Methods**). For HCM this gives us a maximum tolerated allele count of 9, assuming 50% penetrance (or 5 for fully penetrant alleles), for variants genotyped in the full ExAC cohort (sample size=121,412 chromosomes). The MYBPC3:c.1504C>T variant is observed 3 times in ExAC (freq=2.49x10^-5^; Table 1).

To facilitate these calculations, we have produced an online calculator (https://jamesware.shinyapps.io/alleleFrequencyApp/) that will compute maximum credible population allele frequency and maximum sample allele count for a user-specified genetic architecture, and conversely allow users to dynamically explore what genetic architecture(s) might be most compatible with an observed variant having a causal role in disease.

To assess these thresholds empirically, we explored the ExAC allele frequency spectrum of 1132 distinct autosomal variants identified in 6179 recently published HCM cases referred for diagnostic sequencing, and individually assessed and reported according to international guidelines^12,13^. 477/479 (99.6%) variants reported as ‘Pathogenic’ or ‘Likely Pathogenic’ fell below our threshold (Figure 2), including all variants with a clear excess in cases. 419 of these variants are absent from ExAC. The 2 variants historically classified as ‘Likely Pathogenic’, but prevalent in ExAC in this analysis, were reassessed using contemporary ACMG criteria: there was no strong evidence in support of pathogenicity, and they were reclassified in light of these findings (Supplementary Table 1). This analysis identifies 66/653 (10.1%) VUS that are unlikely to be truly causative for HCM.

**Figure 2.**
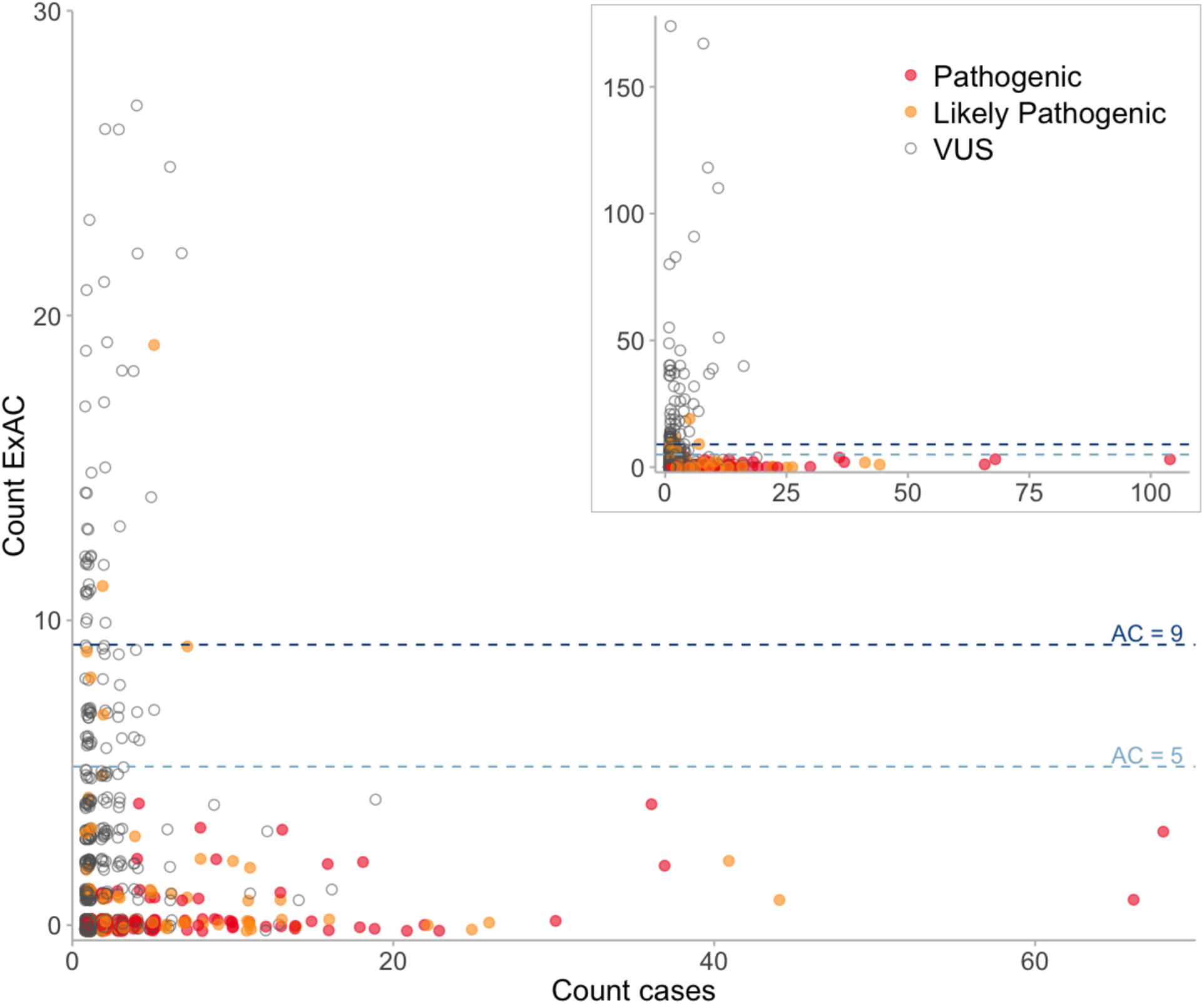
Plot of ExAC allele count (all populations) against case allele count for variants classified as VUS, Likely Pathogenic or Pathogenic in 6179 HCM cases. The dotted lines represent the maximum tolerated ExAC allele counts in HCM for 50 % (dark blue) and 100 % penetrance (light blue). Variants are colour coded according to reported pathogenicity. Where classifications from contributing laboratories were discordant the more conservative classification is plotted. The inset panel shows the full dataset, while the main panel expands the region of primary interest.

The above analysis applied a single global allele count limit of 9 for HCM, however, as allele frequencies differ between populations, filtering based on frequencies in individual populations may provide greater power^2^. For example, a variant relatively common in any one population is unlikely pathogenic, even if rare in other populations, provided the disease prevalence and architecture is consistent across populations. We therefore compute a maximum tolerated AC for each distinct sub-population of our reference sample, and filter based on the highest allele frequency observed in any major continental population (see **Online Methods**).

To further validate this approach, we examined all 601 variants identified in ClinVar^21^ as “Pathogenic” or “Likely Pathogenic” and non-conflicted for HCM. 558 (93%) were sufficiently rare when assessed as described. 43 variants were insufficiently rare in at least one ExAC population, and were therefore re-curated. 42 of these had no segregation or functional data sufficient to demonstrate pathogenicity in the heterozygous state, and we would classify as VUS at most. The remaining variant (MYBPC3:c.3330+5G>C) had convincing evidence of pathogenicity, though with uncertain penetrance (see **Supplementary information**), and was observed twice in the African/African American ExAC population. This fell outside the 95% confidence interval for an underlying population frequency <4x10^-5^, but within the 99% confidence threshold: a single outlier due to stochastic variation is unsurprising given that these nominal probabilities are not corrected for multiple testing across 601 variants. In light of our updated assessment, 20 variants were reclassified as Benign/Likely Benign and 22 as VUS according to the American College for Medical Genetics and Genomics (ACMG) guidelines for variant interpretation^3^ (Supplementary Table 1).

### Extending this approach to other disorders

This approach can be readily applied in diseases where large case series are available to assess the genetic and allelic architecture, such as the inherited cardiac conditions displayed in Table 1. In the absence of large case series, we must estimate the genetic architecture parameters by extrapolating from similar disorders and/or variant databases.

Where disease-specific variant databases exist, we can use these to help estimate the maximum allelic contribution in lieu of individual case series. For example, Marfan syndrome is a rare connective tissue disorder caused by variants in the *FBN1* gene. The UMD-FBN1 database^22^ contains 3077 variants in *FBN1* from 280 references (last updated 28/08/14). The most common variant is in 30/3006 records (1.00%; 95CI 0.53-1.46%), which likely overestimates its contribution to disease if related individuals are not systematically excluded. Taking the upper bound of this frequency as our maximum allelic contribution, we derive a maximum tolerated allele count of 2 (Table 2). None of the five most common variants in the database are present in ExAC.

**Table 2.** Maximum credible population frequencies and maximum tolerated ExAC allele counts for variants causative of exemplar inherited cardiac conditions, assuming a penetrance of 0.5 throughout. CPVT - catecholaminergic polymorphic ventricular tachycardia; FH - familial hypercholesterolaemia. Prevalence estimates were obtained from: Marfan^26^, Noonan^23^, CPVT^24^ and classical Ehlers-Danlos^25^.

| Disease | Maximum allelic contribution | Prevalence | Penetrance* | Maximum population frequency | Maximum tolerated ExAC allele count |
| --- | --- | --- | --- | --- | --- |
| Marfan | 0.015 | 401769 | 0.5 | $5.0 \times 10^{-6}$ | 2 |
| Noonan | 0.100 | 1/1000 | 0.5 | $1.0 \times 10^{-4}$ | 18 |
| CPVT | 0.100 | 1/10,000 | 0.5 | $1.0 \times 10^{-5}$ | 3 |
| Classic Ehlers-Danlos | 0.400 | 1/20,000 | 0.5 | $2.0 \times 10^{-5}$ | 5 |

Where no mutation database exists, we can use what is known about similar disorders to estimate the maximum allelic contribution. For the cardiac conditions with large cases series in Table 1, the maximum proportion of cases attributable to any one variant is 6.7% (95CI 4.1-9.2%; PKP2:c.2146-1G>C found in 24/361 ARVC cases^13^). We therefore take the upper bound of this confidence interval (rounded up to 0.1) as an estimate of the maximum allelic contribution for other genetically heterogeneous cardiac conditions, unless we can find disease-specific evidence to alter it. For Noonan syndrome and Catecholaminergic Polymorphic Ventricular Tachycardia (CPVT - an inherited cardiac arrhythmia syndrome) with prevalences of 1 in 1000^23^ and 1 in 10,000^24^ respectively, this translates to maximum population frequencies of 5x10^-5^ and 5x10^-6^ and maximum tolerated ExAC allele counts of 10 and 2 (Table 2).

Finally, if the allelic heterogeneity of a disorder is not well characterised, it is conservative to assume minimal heterogeneity, so that the contribution of each gene is modelled as attributable to one allele, and the maximum allelic contribution is substituted by the maximum genetic contribution (i.e the maximum proportion of the disease attributable to single gene). For classic Ehlers-Danlos syndrome, up to 40% of the disease is caused by variation in the *COL5A1* gene^25^. Taking 0.4 as our maximum allelic contribution, and a population prevalence of 1/20,000^25^ we derive a maximum tolerated ExAC AC of 5 (Table 2).

Here we have illustrated frequencies analysed at the level of the disease. In some cases this may be further refined by calculating distinct thresholds for individual genes, or even variants. For example, if there is one common founder mutation but no other variants that are recurrent across cases, then it would make sense to have the founder mutation as an exception to the calculated threshold.

### Application to recessive diseases

So far we have considered diseases with a dominant inheritance model. Our framework is readily modified for application in recessive disease, and to illustrate this we consider the example of Primary Ciliary Dyskinesia (PCD), which has a prevalence of up to 1 in 10, 000 individuals in the general population^27^.

Intuitively, if one penetrant recessive variant were to be responsible for all PCD cases, it could have a maximum population frequency of. √(1/10000). The maximum frequency of a recessive disease-causing variant in the population can be more completely defined as:

*max credible allele frequency* = √(*prevalence*) x *maximum allelic contribution* x √(*maximumgeneticcontributiori*)x1/√(*penetrance*)

where *maximum genetic contribution* represents the proportion of all cases that are attributable to the gene under evaluation, and *maximum allelic contribution* represents the proportion of cases attributable to that gene that are attributable to an individual variant (full derivation can be found in **Online Methods**).

We can refine our evaluation of PCD by estimating the maximum genetic and allelic contribution. Across previously published cohorts of PCD cases^28–30^, *DNAI1* IVS1+2_3insT was the most common variant with a total of 17/358 alleles (4.7% 95CI 2.5-7.0%). Given that ~9% of all patients with PCD have disease-causing variants in *DNAI1* and the IVS1+2_3insT variant is estimated to account for ~57% of variant alleles in DNAI1^28^, we can take these values as estimates of the maximum genetic and allelic contribution for PCD, yielding a maximum expected population AF of √ (1/10000)χ 57x √0. 09 x 1/ √0. 5 = 2.42x10^-3^ This translates to a maximum tolerated ExAC AC of 322. *DNAI1* IvSl+2_3insT is itself present at 56/121108 ExAC alleles (45/66636 non-Finnish European alleles). A single variant reported to cause PCD in ClinVar occurs in ExAC with AC > 332 *(NME8* NM_016616.4:c.271-27C>T; AC=2306/120984): our model therefore indicates that this variant frequency is too common to be disease-causing, and consistent with this we note that it meets none of the current ACMG criteria for assertions of pathogenicity, and have reclassified it as VUS (see **Supplementary information**).

### Pre-computing threshold values for the ExAC populations

For each ExAC variant, we defined a “filtering allele frequency” that represents the threshold disease-specific “maximum credible allele frequency” at or below which the disease could not plausibly be caused by that variant. A variant with a filtering allele frequency ≥ the maximum credible allele frequency for the disease under consideration should be filtered, while a variant with a filtering allele frequency below the maximum credible remains a candidate. This value has been pre-computed for all variants in ExAC (see **Online Methods**), and is available via the ExAC VCF and browser (http://exac.broadinstitute.org).

To assess the efficiency of our approach, we calculated the filtering allele frequency based on 60,206 exomes from ExAC and applied these filters to a simulated dominant Mendelian variant discovery analysis on the remaining 500 exomes (see **Online Methods**). Filtering at allele frequencies lower than 0.1% can substantially reduce the number of predicted protein-altering variants in consideration, with the mean number of variants per exome falling from 176 at a cutoff of 0.1% to 63 at a cutoff of 0.0001% (Figure 3a). Additionally, we compared the prevalence of variants in HCM genes in cases and controls across the allele frequency spectrum, and computed disease odds ratios for different frequency bins. The odds ratio for disease-association increases markedly at very low allele frequencies (Figure 3b) demonstrating that increasing the stringency of a frequency filter improves the information content of a genetic result.

**Figure 3.**
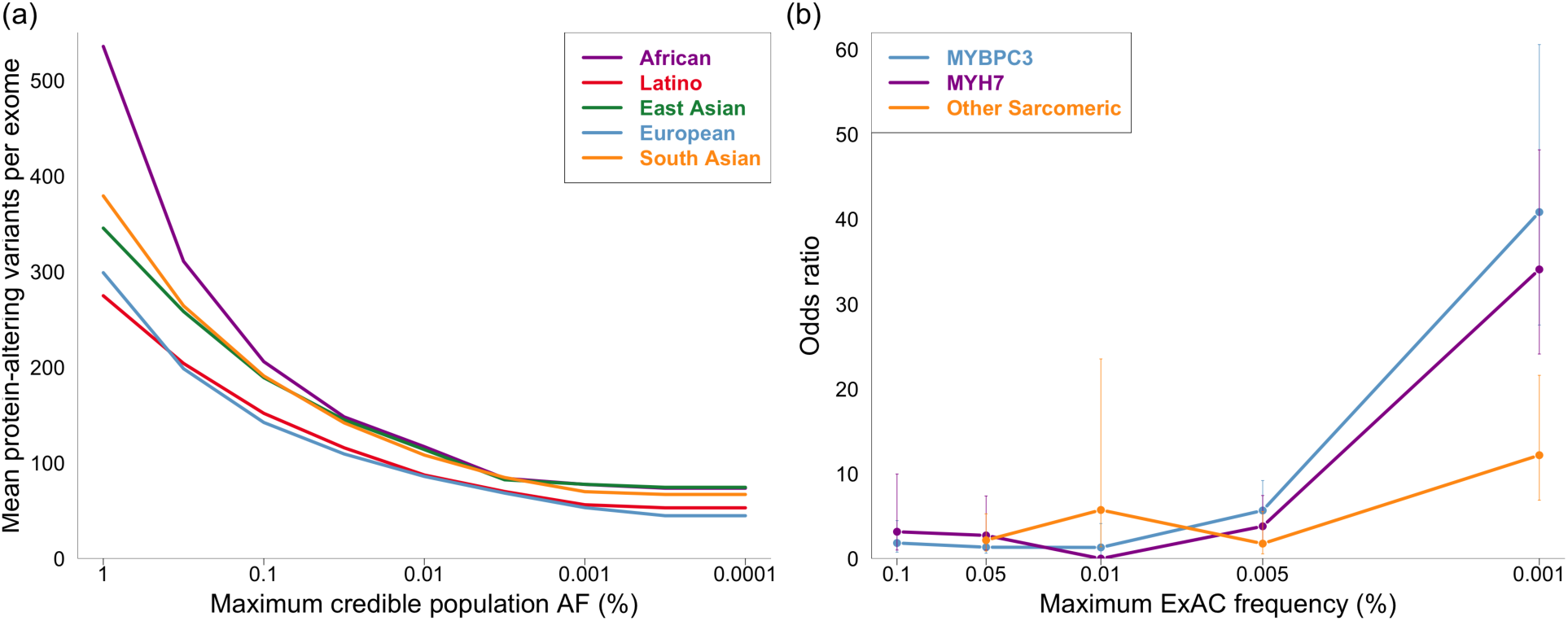
The clinical utility of stringent allele frequency thresholds. (a) The number of predicted protein-altering variants (definition in Online Methods) per exome as a function of the frequency filter applied. A one-tailed 95% confidence interval is used, meaning that variants were removed from consideration if their AC would fall within the top 5 % of the Poisson probability distribution for the user’s maximum credible AF (x axis). (b) The odds ratio for HCM disease-association against allele frequency. The prevalence of variants in HCM-associated genes *(MYH7, MYBPC3* and other sarcomeric *(TNNT2, TNNI3, MYL2, MYL3, TPM1* and *ACTC1,* analysed collectively) in 322 HCM cases and 60,706 ExAC controls were compared for a range of allele frequency bins, and an odds ratio computed (see **Online Methods**). Data for each bin is plotted at the upper allele frequency cutoff. Error bars represent 95 %> confidence intervals. The probability that a variant is pathogenic is much greater at very low allele frequencies.

## DISCUSSION

We have outlined a statistically robust framework for assessing whether a variant is ‘too common’ to be causative for a Mendelian disorder of interest. To our knowledge, there is currently no equivalent guidance on the use of variant frequency information, resulting in inconsistent thresholds across both clinical and research settings. Furthermore, though disease-specific thresholds are recommended^8^, in practice the same thresholds may be used across all diseases, even where they have widely differing genetic architectures and prevalences. We have shown the importance of applying stringent AF thresholds, in that many more variants can be removed from consideration, and the remaining variants have a much higher likelihood of being relevant. We also show, using HCM as an example, how lowering this threshold does not remove true dominant pathogenic variants.

In order to assist others in applying our framework, we have precomputed a ‘filtering allele frequency’ for all variants across the ExAC dataset. This is defined such that if the filtering allele frequency of a variant is at or above the “maximum credible population allele frequency” for the disease in question, then that variant is not a credible candidate (in other words, for any population allele frequency below the threshold value, the probability of the observed allele count in the ExAC sample is <0.05). Once a user has determined their “maximum credible population allele frequency", they may remove from consideration ExAC variants for which the filtering allele frequency is greater than or equal to than the chosen value.

We recognize several limitations of our approach. First, the approach is limited by our understanding of the prevalence and genetic architecture of the disease in question: this characterisation will vary for different diseases and in different populations, though we illustrate approaches to estimation and extrapolation of parameters. In particular, we must be wary of extrapolating to or from less-well characterised populations that could harbour population-specific founder mutations. It is critical to define the genetic architecture in the population under study. Secondly, it is often difficult to obtain accurate penetrance information for reported variants, and it is also difficult to know what degree of penetrance to expect or assume for newly discovered pathogenic variants (see **Supplementary information** for alternative approaches).

Thirdly, while we believe that ExAC is depleted of severe childhood inherited conditions, and not enriched for cardiomyopathies, it could be enriched relative to the general population for some Mendelian conditions, including Mendelian forms of common diseases such as diabetes or coronary disease that have been studied in contributing cohorts. Where this is possible, the maximum credible population allele frequency can be simply computed based on the estimated disease prevalence in the ExAC cohort, rather than the population prevalence. Finally, although the resulting allele frequency thresholds are more stringent than those previously used, they are likely to still be very lenient for many applications. For instance, we base our calculation on the most prevalent known pathogenic variant from a disease cohort. For HCM, for which more than 6,000 people have been sequenced, it is unlikely that any single newly identified variant, not previously catalogued in this large cohort, will explain a similarly large proportion of the disease as the most common causal variant, at least in well-studied populations. Future work may therefore involve modeling the frequency distribution of all known variants for a disorder, to further refine these thresholds.

The power of our approach is limited by currently available datasets. Increases in both the ancestral diversity and size of reference datasets will bring additional power to our method over time. We have avoided filtering on variants observed only once, because a single observation provides little information about true allele frequency. A ten-fold increase in sample size, resulting from projects such as the US Precision Medicine Initiative, will separate vanishingly rare variants from those whose frequency really is ~1 in 100,000. Increased phenotypic information linked to reference datasets will also reduce limitations due to uncertain disease status, and improve prevalence estimates, adding further power to our approach.

## ACKNOWLEDGEMENTS

This work was supported by the Wellcome Trust (107469/Z/15/Z), the Medical Research Council (UK), the NIHR Biomedical Research Unit in Cardiovascular Disease at Royal Brompton & Harefield NHS Foundation Trust and Imperial College London, the Fondation Leducq (11 CVD-01), a Health Innovation Challenge Fund (HICF-R6-373) award from the Wellcome Trust and Department of Health, UK, and by the National Institute of Diabetes and Digestive and Kidney Diseases and the National Institute of General Medical Sciences of the NIH (awards U54DK105566 and R01GM104371). EVM is supported by the National Institutes of Health under a Ruth L. Kirschstein National Research Service Award (NRSA) NIH Individual Predoctoral Fellowship (F31) (award AI122592-01A1). AHO-L is supported by National Institutes of Health under Ruth L. Kirschstein National Research Service Award 4T32GM007748.

This publication includes independent research commissioned by the Health Innovation Challenge Fund (HICF), a parallel funding partnership between the Department of Health and Wellcome Trust. The views expressed in this work are those of the authors and not necessarily those of the Department of Health or Wellcome Trust.

## DATA AVAILABILITY

All data required to reproduce these analyses is available at https://github.com/ImperialCardioGenetics/frequencyFilter. The manuscript was compiled in R, and source code for the analysis, figures and manuscript, are available at the same location. Curated variant interpretations are deposited in ClinVar *Accession & DOI to be added.* ExAC annotations are available at. Our allele frequency calculator app is located at https://jamesware.shinyapps.io/alleleFrequencyApp/, and the source code available at http://github.com/jamesware/alleleFrequencyApp.

## ONLINE METHODS

### Calculating maximum tolerated allele counts

The maximum frequency of a dominant disease-causing variant in the population was defined as:

*maximum credible population AF = prevalence × maximum allelic contribution* × *1/penetrance*

Estimates of disease prevalence were obtained from the literature. Where multiple different values were reported, the highest was used in the calculation, which leads to lenient filtering. A variant penetrance of 0.5 was used for all analyses, as penetrance estimates for individual variants are not widely available. This corresponds to the reported penetrance of the HCM variant used to illustrate our approach^14^ and is the minimum found when researching other variants/disorders.

Determination of the maximum allelic contribution (a measure of heterogeneity) is described in the text. Where a large cohort exists for a disorder, the upper confidence interval of the frequency of the most common variant in this cohort, was used as the maximum allelic contribution.

Having established a maximum credible allele frequency (AF), the maximum tolerated allele count (AC) was computed as the AC occurring at the upper bound of the one-tailed 95% confidence interval (95%CI AC) for that allele frequency, given the observed allele number (AN). Since the population is drawn without replacement, this would strictly be a hypergeometric distribution, but this can be modeled as binomial as the sample is much smaller than the population from which it is drawn. For ease of computation, we approximate this with a Poisson distribution. In R, this is implemented as `max_ac = qpois(quantile_limit,an*af)`, where max_ac is the 95%CI AC, quantile_limit is 0.95 (for a one-sided 95%CI), an is the observed allele number, and af is the maximum credible population allele frequency.

### Application to recessive diseases

The prevalence of a recessive condition can be related to the allele frequency of causative variants by:

*Prevalence =* Σ *(allele frequency of causative alleles in each contributing gene)*^2^ × *penetrance*

approximating to:

*Prevalence = (combined frequency of causative alleles in gene)^2^ × (number of similar genes) x penetrance* and expanding to:

*Prevalence = (max individual allele frequency × 1/maximum allelic contribution)^2^ × 1/maximum genetic contribution × penetrance*

where *maximum genetic contribution* represents the proportion of all cases that are attributable to the gene under evaluation, and *maximum allelic contribution* represents the proportion of cases attributable to that gene that are attributable to an individual variant. The maximum frequency of a recessive disease causing variant in the population was therefore defined as:

*max credible allele frequency* = √*(prevalence) × maximum allelic contribution ×* √ *(maximumgeneticcontribution) × 1/*√*(penetrance)*

### Pre-computing filtering allele frequency values for ExAC

We define the “filtering allele frequency” for a variant, or `af_filter`, as the highest true population allele frequency for which the upper bound of the 95% confidence interval of allele count under a Poisson distribution is still less than the variant’s observed allele count in the reference sample. It functions as equivalent to a lower bound estimate for the true allele frequency of an observed variant: if the filtering allele frequency of a variant is at or above the maximum credible allele frequency for a disease, then the variant is considered too common to be causative of the disease.

Consider, for example, a variant with an observed AC=3 and AN=100,000. If a user’s maximum credible allele frequency for their disease is 1 in 100,000, then this variant should be kept in consideration as potentially pathogenic, because the upper bound of the Poisson 95%CI is AC=3. On the other hand, if the user’s credible tolerated allele frequency is 1 in 200,000 then this variant should be filtered out, as the 95%CI upper bound is only AC=2. We define `af_filter` as the highest AF value for which a variant should be filtered out.

In the example, the highest allele frequency that gives a 95%CI AC of 2 when AN=100,000 is approximately 8.17e-6. Instead of solving exactly for such values, which would require solving the inverse cumulative distribution function of the Poisson distribution, we derive a numerical approximation in two steps:

1. For each variant in consideration, we use R’s uniroot function to find an AF value (though not necessarily the highest AF value) for which the 95%CI AC is one less than the observed AC.
2. We then loop, incrementing by units of millionths, and return the highest AF value that still gives a 95%CI AC less than the observed AC.

In order to pre-compute `af_filter` values for all of ExAC (verson 0.3.1), we apply this procedure to the AC and AN values for each of the five major continental populations in ExAC, and take the highest result from any population. Usually, this is from the population with the highest nominal allele frequency. However, because the tightness of a 95% confidence interval in the Poisson distribution depends upon sample size, the stringency of the filter depends upon the allele number (AN). The stringency of the filter therefore varies appropriately according the the size of the sub-population in which the variant is observed, and sequencing coverage at that site, and `af_filter` is occasionally derived from a population other than the one with the highest nominal allele frequency.

For this analysis, we used adjusted AC and AN, meaning variant calls with GQ≥20 and DP≥10.

### Treatment of singletons and other populations

It is worth considering whether a single observation in a reference sample should ever be treated as incompatible with disease. Using the approach outlined above, it can be inferred that an ExAC AC=1 would be considered incompatible with a true population allele frequency <2.9x10^-6^ (with 95% confidence). For a penetrant disease with a prevalence of 1:1,000,000, the probability of observing a specific causative allele in ExAC is <0.01, even if the disease is genetically homogeneous with just one causative variant. In practice however, we feel that there are few, if any, diseases that are extremely rare yet have sufficiently well-characterized genetic architecture to discard singleton variants from a reference sample. Therefore, for singletons (variants observed exactly once in ExAC), we set the filtering allele frequency to zero.

We also note that occasionally a variant is seen in individuals falling under the Finnish or “Other” population categories in ExAC, and is a singleton or absent in all five continental populations. For these variants, the filtering allele frequency is set to zero. Because the Finnish are a bottlenecked population, disease-causing alleles may reach frequencies that would be impossible in large outbred populations. Similarly, because we have not assigned ancestry for the “Other” individuals, it is difficult to assess the population frequency of variants seen only in this set of individuals. Users are left to judge whether variants that would not be filtered on the basis of frequency in the five continental populations, but that are recurrent in Finnish or “Other” populations, should be removed from consideration according to the specific circumstances.

### Simulated Mendelian variant discovery analysis

To simulate Mendelian variant discovery, we randomly selected 100 individuals from each of five major continental populations and filtered their exomes against filtering allele frequencies derived from the remaining 60,206 ExAC individuals. The subset of individuals was the same as that previously reported^2^. Predicted protein-altering variants are defined as missense and equivalent (including in-frame indels, start lost, stop lost, and mature miRNA-altering), and protein-truncating variants (nonsense, essential splice site, and frameshift).

### Variant curation

We utilized the July 9, 2015 release of ClinVar, extracting variants from XML and TXT releases into a single tab-delimited file through use of a Python implementation of vt normalize^31^, as described previously^2^. Only variants annotated as pathogenic and non-conflicted were investigated. ExAC counts were determined by matching on chromosome, position, reference, and alternate alleles. For all variants above the proposed maximum tolerated allele count for HCM, all HGMD annotated literature was reviewed and the level of evidence supporting disease pathogenicity was curated according to ACMG criteria^3^.

### Calculating odds ratios for HCM variant burden

We used a cohort of 322 patients recruited to the Royal Brompton Hospital cardiac Biomedical Research Unit with diagnosis of HCM confirmed by cardiac MRI. These samples were sequenced using the IlluminaTruSight Cardio Sequencing Kit^32^ on theIlluminaMiSeq and NextSeq platforms. This study was subject to ethical approval (REC: 09/H0504/104+5) and informed consent was obtained for all subjects. The number of rare variants in *MYBPC3, MYH7*and the six other sarcomeric genes associated with HCM ( *TNNT2, TNNI3, MYL2, MYL3, TPM1* and *ACTC1)* were calculated for this HCM cohort, and for reference population samples from ExAC. Case/control variant frequencies were calculated for all protein altering variants (frameshift, nonsense, splice donor/acceptor, missense and in-frame insertions/deletions), with frequencies and case/control odds ratios calculated separately for non-overlapping ExAC allele frequency bins with the following breakpoints: 1x10^-5^, 5x10^-5^, 1x10^-4^, 5x10^-4^ and 1x10^-3^. Odds Ratios were calculated as OR = (cases with variant / cases without variant) / (ExAC samples with variant / ExAC samples without variant) along with 95% confidence intervals. In the absence of sample-level genotype data for ExAC, the number of samples with a variant was approximated by the total number of variant alleles - i.e. assuming that each rare variant was found in a distinct sample.

## CODE AVAILABILITY

The manuscript was compiled in R, and source code for the analysis, figures and manuscript, are available at https://github.com/ImperialCardioGenetics/frequencyFilter. The source code for our allele frequency calculator app is located at http://github.com/jamesware/alleleFrequencyApp.

## SUPPLEMENTARY INFORMATION

Supplementary note 1 - Curation of a high frequency PCD variant

Supplementary note 2 - Dealing with penetrance

Supplementary table 1

